# Reference genes for mesangial cell and podocyte qPCR gene expression studies under high-glucose and renin-angiotensin-system blocker conditions

**DOI:** 10.1101/2021.01.19.427251

**Authors:** Nicole Dittrich Hosni, Ana Carolina Anauate, Mirian Aparecida Boim

**Affiliations:** Nephrology Division, Department of Medicine, Universidade Federal de São Paulo, São Paulo, Brazil

**Keywords:** Essential genes, Diabetic Nephropathy, Mesangial Cells, Podocytes, Real-Time Polymerase Chain Reaction, Renin-Angiotensin System, In Vitro Techniques

## Abstract

**Background:** Real-time PCR remains currently the gold standard method for gene expression studies. Identification of the best reference gene is a key point in performing high quality qPCR, providing strong support for results, as well as performing as a source of bias when inappropriately chosen. Mesangial cells and podocytes, as essential cell lines to study diabetic kidney disease (DKD) physiopathology, demand accurate analysis of the reference genes used so far to enhance validity of gene expression studies, especially regarding high glucose (HG) and DKD treatments, with angiotensin II receptor blockers (e.g. Losartan) being the most commonly used. This study aimed to evaluate the suitability and define the most stable reference gene for mesangial cells and podocytes studies of an *in vitro* DKD model of disease and its treatment.

**Methods:** Five software packages (RefFinder, NormFinder, GeNorm, Bestkeeper, and DataAssist) and the comparative ΔCt method were selected to analyze six different candidate genes: *HPRT, ACTB, PGAM-1, GAPDH, PPIA,* and *B2M.* RNA was extracted and cDNA was synthesized from immortalized mouse mesangial cells and podocytes cultured in 4 groups: control (n=5; 5mM glucose), mannitol (n=5; 30mM, as osmotic control), HG (n=5; 30mM glucose), and HG + losartan (n=5; 30mM glucose and 10^-4^ mM of losartan). Real-time PCR was performed according to MIQE guidelines.

**Results:** We identified that the use of 2 genes is the best combination for qPCR normalization for both mesangial cell and podocytes. For mesangial cells, the combination of *HPRT* and *ACTB* presented higher stability values. For podocytes, *HPRT* and *GAPDH* showed the best results.

**Conclusion:** This analysis provides support for the use of *HPRT* and *ACTB* as reference genes in mouse mesangial cell studies of gene expression via real-time PCR technique, while for podocytes, *HPRT* and *GAPDH* should be chosen.

## INTRODUCTION

Globally, diabetic kidney disease (DKD) related deaths are increasing compared to other types of chronic kidney diseases (1). Diabetes endures as the dominant cause of end-stage renal disease, being responsible for approximately half of cases in developed countries (2).

DKD development triggers glomerular injuries, including hyperfiltration, progressive albuminuria, declining glomerular filtration rate, and eventually end-stage renal disease (3). Additionally, early cellular damage appears in mesangial cells and podocytes (4). Characteristic features of mesangial damage relies on mesangial expansion, following cell enlargement, secretion of extracellular matrix, and ultimately nodular glomerulosclerosis (5). Commonly, podocytes exposed to a high glucose environment develop foot process effacement, hypertrophy, detachment from the basal membrane, and apoptosis (6,7).

Analysis of gene expressions *in vivo* and *in vitro* models of DKD are among the strategies that contribute to better understand the pathophysiological mechanisms of progression of DKD. The very sensitive quantitative real-time PCR (qPCR) is currently the gold standard method to evaluate gene expression (8). Identification of the best reference gene stands as a key point in performing high-quality qPCR, providing strong support for results, as well as acting as a source of bias when inappropriately chosen. Considering the many steps the procedure goes through (RNA extraction, reverse-transcription, amplification efficiency, etc.) and the fact that the data is most frequently relative, not absolute, normalization is established as a critical step to properly standardize the experiment and, thus, provide decisive results for a qPCR assay. Although the use of reference genes is absolutely acknowledged as the most correct method of normalization, gene choice must be validated according to tissue, cell type, experimental design, and conditions (9). There must be a detailed report of the method used to select the most stable gene and the optimal number of genes recommended (10). Specific celltypes from glomeruli, however, do not have support from the literature concerning the best normalization gene for qPCR studies, circumstance that may complicate the interpretation of qPCR data for researchers on the field, misrepresenting the results’ reliability.

Mesangial cells and podocytes, as essential cell lines in DKD, demand accurate analysis of the most excellent reference genes to enhance validity of gene expression studies in the field, especially regarding high glucose (HG) and different treatments, being angiotensin II receptor blockers the most frequently used (11,12). Unfortunately, to our knowledge, the only available reference in the literature regarding DKD qPCR reference genes relies on entire glomerulus analysis, not suitable to cell specific *in vitro* assays (13).

Our goal was to evaluate the suitability and define the most stable reference gene specifically for mesangial cell and podocyte studies of an *in vitro* DKD model of disease and its treatment, among six commonly used reference genes *(HPRT, ACTB, PGAM-1, GAPDH, PPIA,* and *B2M).*

## MATERIALS AND METHODS

### Cell lines and cell culture

Immortalized mesangial cells (SV40 MES 13, ATCC) were cultured in DMEM (Invitrogen Corporation, Gaithersburg, MD, USA) containing 10% fetal bovine serum (FBS), penicillin (50 U/ml) and 2.6 g HEPES at 37°C. Podocytes (Cell line E11, CLS) were cultured in RPMI 1640 medium (Invitrogen Corporation, Gaithersburg, MD, USA) supplemented with 10% fetal bovine serum and interferon-gama *(INF-gama)* at 33 °C; after achieving the desired confluence, flasks were transferred to a 37 °C incubator for the differentiation process for 14 days without *INF*-gama. Both cell types were cultured until >90% of confluence and remained in a 5% CO_2_ environment. After 24 hours in 1% FBS, each group received the designated stimulus for 24 hours: pure medium (control group), medium containing 30 mM mannitol (as osmotic control, mannitol group), or 30mM D-glucose (high-glucose group) or 30mM D-glucose combined with 100uM losartan (losartan group). The study workflow is shown in Figure 1.

**Fig. 1.**
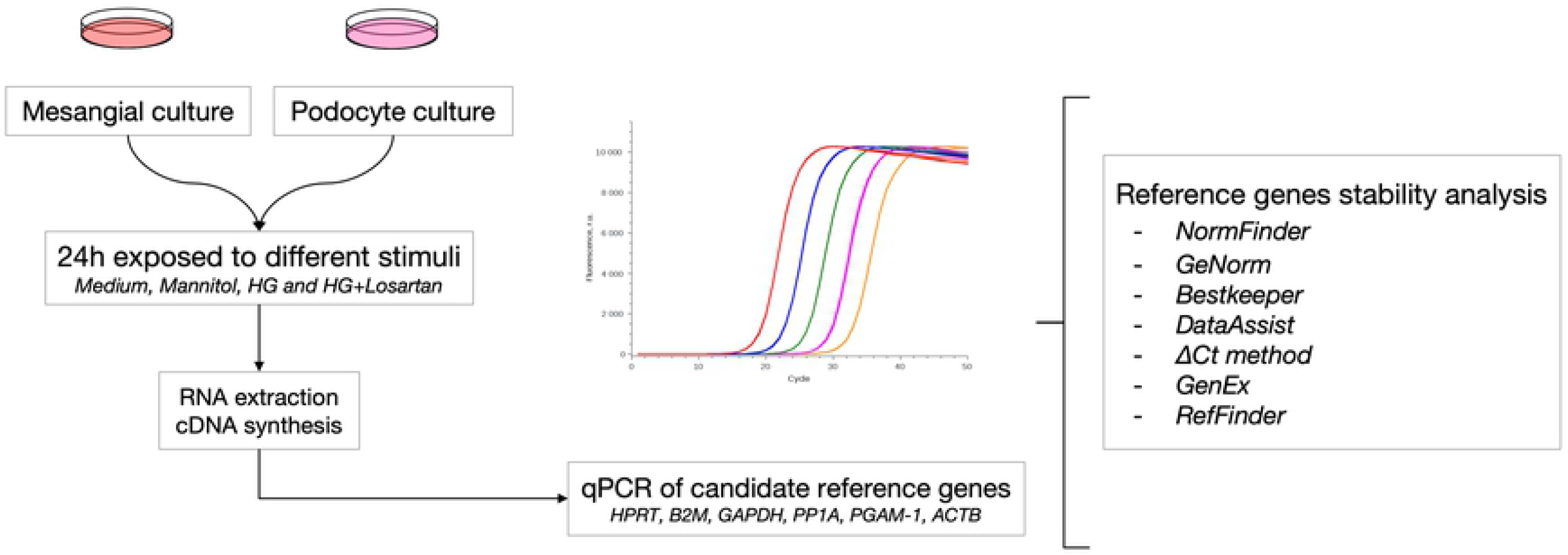
Study workflow. The figure shows the workflow for determination of the most stable reference gene for mesangial cells and podocytes exposed to mannitol, high glucose or high glucose and losartan.

### RNA extraction, quality parameters and reverse transcription

Total RNA was extracted using TRIzol reagent (Life Technologies, USA), according to the manufacturer’s instructions. RNA concentration and quality (260/280 ratio >1.8 and 260/230 ratio 2.0-2.2, indicating high purity) was assessed using the NanoVue spectrophotometer (GE Healthcare Life Sciences, USA). RNA integrity was also analyzed by gel electrophoresis. After RNA extraction, we performed DNAse treatment to avoid genomic DNA contamination. An amount of 2 μg of total RNA was reverse-transcribed into cDNA (High Capacity cDNA Reverse Transcription Kit, Applied Biosystems, USA). The reaction mixture was incubated for 10 minutes at 25 °C, 120 minutes at 37 °C and 5 seconds at 85 °C.

### qPCR performance

Gene expression analysis was performed by qPCR using SYBR Green (Applied Biosystems) in QuantStudio 7 Flex (Applied Biosystems), in accordance with the manufacturer’s instructions. Primer sequences for the six genes used are exposed in Table 1. Melting curves of all primers are exposed in Figure 2. All samples were evaluated in triplicate.

**Fig. 2.**
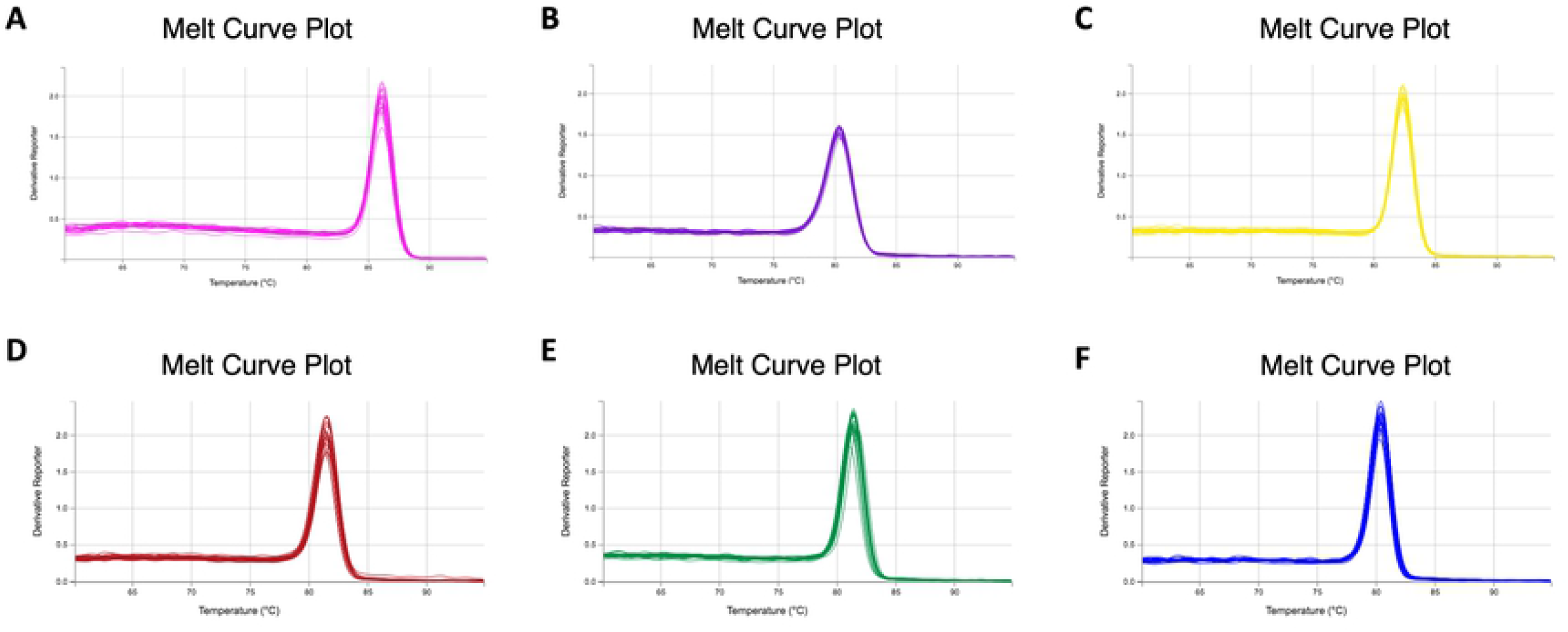
Melting curves of the primers for the candidate genes. A) ACTB. B) B2M. C) PGAM- 1. D) GAPDH. E) HPRT. F) PPIA.

**Table 1.**
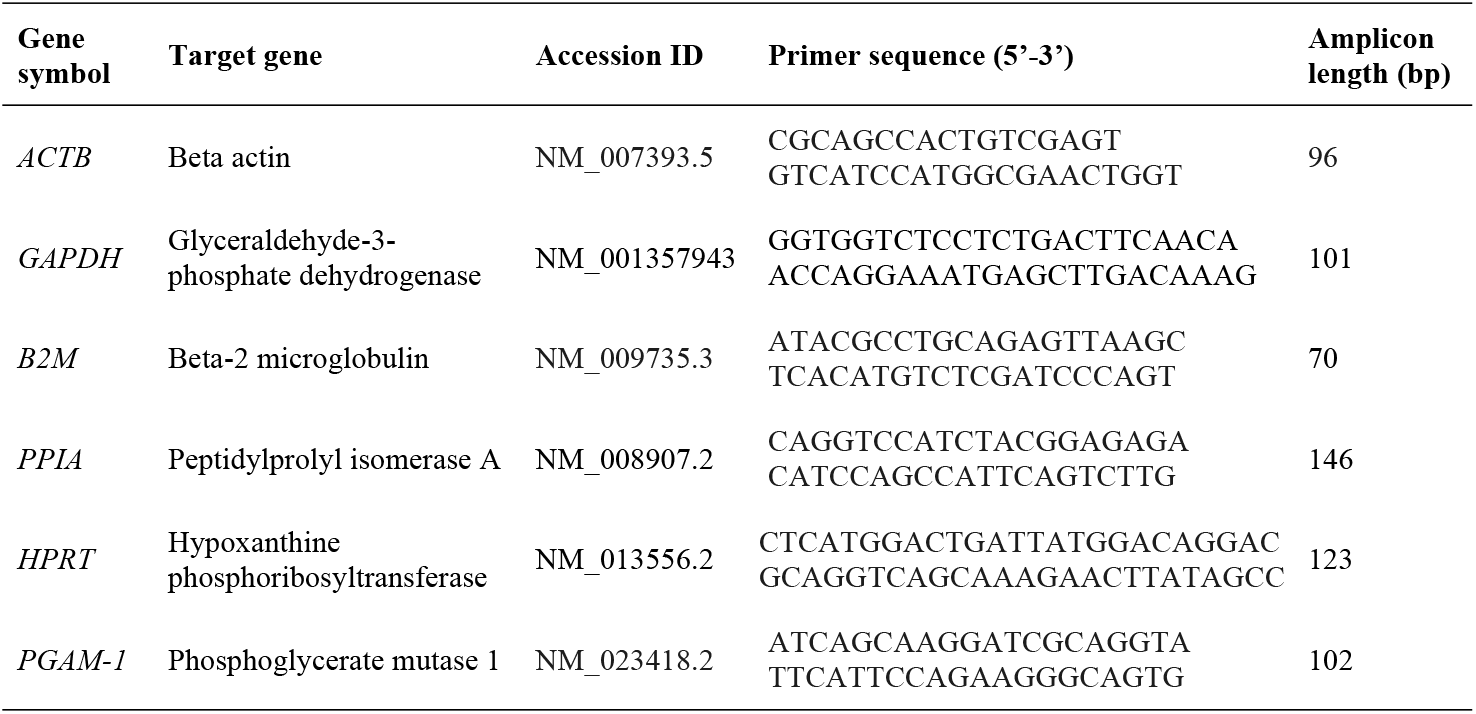
Primer sequences for the six candidate genes.

### Software analysis for stability of candidate reference genes

To establish the best reference gene and best combination, we evaluated qPCR results in five different software applications: RefFinder, NormFinder, GeNorm, Bestkeeper, and DataAssist. We also evaluated the data with the comparative ΔCt method.

NormFinder is a freely available tool that provides the stability value for several candidate genes tested on a sample set. Any required number of samples are subject to the analysis, providing an estimation of expression variation (14). GeNorm software works as an algorithm (M value) to determine the most stable reference genes among a collection of tested candidate genes. The tool calculates a normalization factor for each sample, established according to the geometric mean of the reference genes number (10). Bestkeeper is an excel based spreadsheet software that determines the best suited reference genes and combines them into an index, allowing a comparison with further target genes to decide which of them has the best suitability for normalization. The application acknowledges extremely deviating samples, that can be removed from the calculation and improve the results reliability (15). DataAssist is an Applied Biosystems’ software that quantifies relative gene expression across a given number of samples. It provides an “Endogenous Control Selection” tool that shows the Ct values of candidate genes for all samples as well as a score (16).The ΔCt method compares the relative expression values between ‘pairs of genes’, implementing an elimination process according to a ranking of the variability among each pair. Subsequently, the most appropriate gene of reference can be selected (17).

The program GenEx was used in order to calculate the accumulated standard deviation across the samples, providing the necessary number of genes required for the minimum standard deviation (18). Finally, we used the RefFinder software, an all-encompassing program developed with the aim to evaluate reference genes from experimental data. The tool includes available software algorithms and methods, all of them previously mentioned: geNorm, Normfinder, BestKeeper, and the comparative ACt method. Supported by the ranking of each program, the RefFinder calculates the geometric mean for an overall final ranking (19).

### Statistical Analysis

The entire dataset was analyzed regarding normality (Shapiro-Wilk test) and homogeneity (Levene’s test). All comparisons were analyzed using ANOVA or Kruskal-Wallis, according to each test prerequisites. The level of significance considered was p<0.05. Analysis was performed on Jamovi software, version 1.0.1. Results are expressed as mean ± standard deviation (SD).

## RESULTS

### Expression levels profile of candidate genes of reference

Raw Ct values were acquired in triplicate for both mesangial cell and podocytes samples and analyzed according to each stimulus received. Ct values are inversely proportional to the gene expression. The Ct mean of the candidate genes ranged from 29.20 to 18.55 in mesangial cells. The highest Ct among the candidate genes in mesangial cells was achieved by *ACTB* (29.20 ± 1.09), and the lowest by *PPIA* (18.55 ± 0.79). *HPRT* showed a mean of 23.48 ± 0.97, followed by *GAPDH* (22.46 ± 1.04), *PGAM-1* 21.86 ± 1.06 and *B2M* 18.66 ± 0.78.

For podocytes, otherwise, the mean ranged from 24.45 to 13.02. *ACTB* achieved the highest value (24.45 ± 1.2), while the lowest value was achieved by *B2M* (13.02 ± 0.51). The remaining candidates showed a mean between 19.19 and 14.98: *GAPDH* (19.19 ± 1.00) was followed by *HPRT*(18.85 ± 0.79), *PGAM-1* (17.90 ± 1.16) and *PPIA* (14.98 ± 1.06). The mean Ct value of the triplicates according to each gene and cell line is shown in Figures 3A and 3B.

**Fig. 3.**
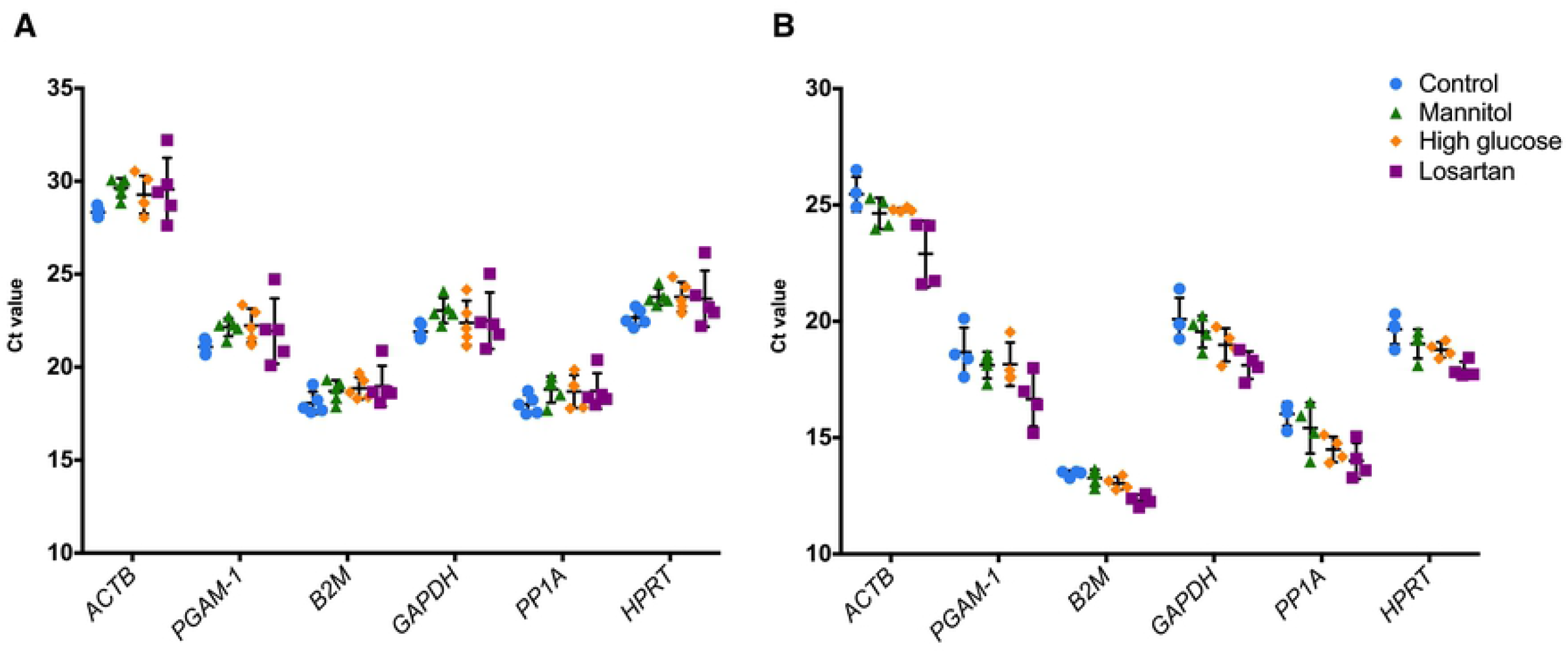
Expression profile of the six candidate reference genes in mesangial cells (A) and podocytes (B). A lower threshold value (Ct) represents a higher gene expression level. The data are presented as mean +/- standard deviation. Each dot represents the average from triplicated ΔCt from each sample. All genes were tested for differences among the groups (Kruskal-Wallis test) and all comparisons showed a non-significant result (p>0.05).

### Stability of candidate genes

We applied the six algorithms described previously to determine the stability of each reference gene candidate according to the cell type. After analysis with different algorithms and a visual inspection of the ranked genes, we concluded that *HPRT* and *ACTB* for mesangial cells (Table 2) and *HPRT* together with *GAPDH* for podocytes were the best reference genes for qPCR studies (Table 3).

NormFinder showed the lowest stability value for *HPRT* for both mesangial cells and podocytes, determining this gene as the best reference gene according to this algorithm. As recommended by software instructions, any gene with a stability value higher than 0.5 is considered unstable - all genes tested showed a stability value lower than the cutoff.

Individual results for BestKeeper showed, for mesangial cells, the lowest coefficient of variation (CV) for *ACTB* thus, indicating this gene as the best for this cell type under the established conditions according to this software. For podocytes, the lowest CV was displayed for *B2M,* showing up as the most stable option. No candidate gene showed a SD higher than 1.0, the fixed software threshold for instability.

The DataAssist software retrieved as the most stable gene for mesangial cells the *PGAM-1* gene. Nevertheless, the lowest score for podocytes was achieved by *HPRT.*

Regarding the ΔCt method, the lowest SD was obtained by *HPRT* for both cell lines. The highest SD was shown by *B2M* for mesangial cells and *ACTB* for podocytes, classifying them as the least stable genes according to the method.

For mesangial cells, GeNorm showed the best results for *ACTB* and *GAPDH* together according to M-values. *B2M* was considered the least stable gene. For podocytes, the best pair of M-values was given to *HPRT* and *GAPDH.* The least stable gene for podocytes was *PPIA.*

Based on these results and visual inspection of all data (Table 2 and 3), *HPRT* was selected as the overall best reference gene for both mesangial cells and podocytes. For mesangial cells, *HPRT* and *ACTB* were considered the best combination of genes for qPCR normalization (Table 2). *PPIA,* otherwise, was classified as the least stable for mesangial cells. Along with *HPRT*, *GAPDH* was also ranked as the most stable candidate reference gene for podocytes, while *ACTB, PGAM-1,* and *PPIA* were found the least feasible genes (Table 3).

**Table 2.** Ranking of candidate reference genes by each method used for mesangial cells.

| NormFinder* | Stability value | GeNorm | M value | BestKeeper | CV [% CP] | std dev [± CP] | DataAssist | Score | RefFinder | Geomean | ΔCt method | Mean SD | Visual inspection** | Frequency |
| --- | --- | --- | --- | --- | --- | --- | --- | --- | --- | --- | --- | --- | --- | --- |
| *HPRT* | 0.118 | *ACTB* | 0.031 | *ACTB* | 2.96 | 0.87 | *PGAM-1* | 5.324 | *HPRT* | 1.00 | *HPRT* | 0.67 | *HPRT* | 3x |
| *ACTB* | 0.158 | *GAPDH* | 0.031 | *HPRT* | 2.99 | 0.70 | *PPIA* | 5.514 | *ACTB* | 2.21 | *ACTB* | 0.75 | *ACTB* | 2x |
| *GAPDH* | 0.179 | *PGAM-1* | 0.034 | *B2M* | 3.03 | 0.57 | *HPRT* | 5.525 | *PPIA* | 3.41 | *PGAM-1* | 0.77 | *PGAM-1* | 1x |
| *PGAM-1* | 0.181 | *HPRT* | 0.038 | *PPIA* | 3.41 | 0.63 | *ACTB* | 5.624 | *PGAM-1* | 3.94 | *PPIA* | 0.77 | *GAPDH* | 1x |
| *PPIA* | 0.226 | *PPIA* | 0.039 | *GAPDH* | 3.51 | 0.79 | *GAPDH* | 5.907 | *GAPDH* | 4.47 | *GAPDH* | 0.86 | *B2M* | 0x |
| *B2M* | 0.244 | *B2M* | 0.042 | *PGAM-1* | 3.65 | 0.80 | *B2M* | 6.835 | *B2M* | 4.56 | *B2M* | 0.92 | *PPIA* | 0x |

Lower values indicate increased stability in gene expression. Each software result is shown in order of stability.
\*Best reference genes determined by NormFinder when the intra- and intergroup variations were not considered.
\*\*Visual inspection refers to the number of times each gene appears as the top one in each analysis

**Table 3.** Ranking of candidate reference genes by each method used for podocytes.

| NormFinder* | Stability value | GeNorm | M value | BestKeeper | CV [% CP] | std dev [± CP] | DataAssist | Score | RefFinder | Geomean | ΔCt method | Mean SD | Visual inspection** | Frequency |
| --- | --- | --- | --- | --- | --- | --- | --- | --- | --- | --- | --- | --- | --- | --- |
| *HPRT* | 0.174 | *HPRT* | 0.030 | *B2M* | 0.43 | 13.64 | *HPRT* | 0.54 | *HPRT* | 1.32 | *HPRT* | 0.66 | *HPRT* | 5x |
| *GAPDH* | 0.204 | *GAPDH* | 0.030 | *HPRT* | 0.65 | 20.31 | *GAPDH* | 0.56 | *GAPDH* | 2.21 | *GAPDH* | 0.69 | *B2M* | 1x |
| *PPIA* | 0.316 | *B2M* | 0.037 | *GAPDH* | 0.79 | 21.40 | *PGAM-1* | 0.66 | *PGAM-1* | 2.59 | *PGAM-1* | 0.79 | *GAPDH* | 1x |
| *PGAM-1* | 0.324 | *ACTB* | 0.044 | *PGAM-1* | 0.82 | 20.13 | *PPIA* | 0.69 | *B2M* | 2.83 | *B2M* | 0.88 | *PGAM-1* | 0x |
| *B2M* | 0.349 | *PGAM-1* | 0.049 | *ACTB* | 0.87 | 26.50 | *B2M* | 0.72 | *PPIA* | 5.23 | *PPIA* | 0.90 | *PPIA* | 0x |
| *ACTB* | 0.373 | *PPIA* | 0.053 | *PPIA* | 0.88 | 16.51 | *ACTB* | 0.85 | *ACTB* | 5.42 | *ACTB* | 1.01 | *ACTB* | 0x |

Lower values indicate increased stability in gene expression. Each software result is shown in order of stability.
\*Best reference genes determined by NormFinder when the intra- and intergroup variations were not considered.
\*\*Visual inspection refers to the number of times each gene appears as the top one in each analysis.

### Determination of the suitable number of reference genes

For each cell line, we determined the optimal number of genes to be used in a gene expression experiment via qPCR. This analysis was performed by the Genex software and the accumulated standard deviation (Acc.SD) parameter was considered for each cell line according to the number of genes used. For mesangial cells, we concluded that the Acc.SD decreased proportionally to the number of genes used. We also observed that the difference from one to two genes was higher than 0.1. However, the difference from two to three genes was smaller than 0.1 - a pattern that could be noticed in the following number of genes as well, achieving a *plateau*. Therefore, it would be reasonable to use two reference genes (*HPRT* and *ACTB}* and maintain a smaller source of error, since a higher number of genes increases the overall noise of the experiment as well as the cost (Figure 4A).

**Fig. 4.**
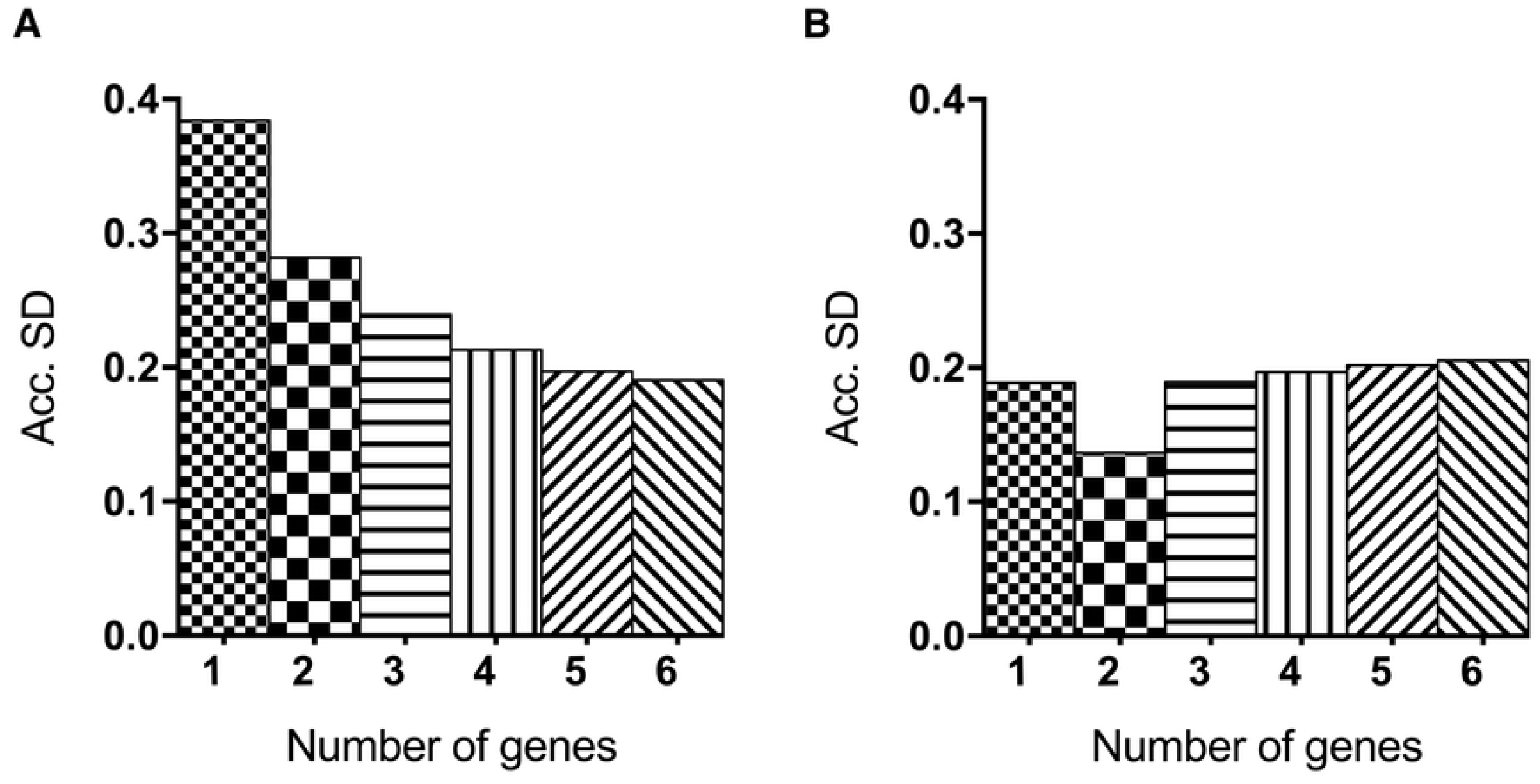
Evaluation of the optimal number of reference genes in (A) mesangial cells and (B) podocyte cells. Accumulated standard deviation (Acc.SD) was accessed by the GenEx software for the six candidate reference genes in all samples for each cell type. Lower values of Acc.SD indicate the best number of reference genes.

For podocytes, the lowest Acc.SD was acquired in the presence of 2 reference genes (Figure 4B). In this case, the use of 2 genes - *HPRT* and *GAPDH or B2M*, the top genes according to the visual inspection - to analyze qPCR results would be the best option as well. Since *GAPDH* and *B2M* showed the same results on visual inspection ranking, we looked closely to the performance of each gene on all softwares: besides being the top gene for BestKeeper, *B2M* appeared in 5^th^ place for NormFinder, 3^rd^ for GeNorm, 5^th^ for DataAssist, 4^th^ for RefFinder and 4^th^ again for ΔCt method. *GAPDH*, however, in addition to being the top gene in GeNorm, appeared as 2^nd^ for NormFinder, 3^rd^ for BestKeeper, 2^nd^ for DataAssist, 2^nd^ for RefFinder and 2^nd^ for the ΔCt method. Considering the overall performance of both genes, *GAPDH* was selected as a best option to pair with *HPRT* as reference genes for podocytes.

### Correlation between the top candidates

After determining that the use of 2 reference genes would be the ideal option for mesangial cells and podocytes, we checked if the best 2 genes for each cell line were correlated and therefore could be used simultaneously. We found that the 2 recommended genes for mesangial cells, *HPRT* and *ACTB,* were strongly correlated (*ρ*=0.80, p<0.0001, Figure 5A), providing support to the recommendation of using these genes at the same time to analyze qPCR data. The same occurred for podocytes: there was a strong correlation between *HPRT* and *GAPDH* expression data, again supporting the use of those genes together as reference genes for qPCR (*ρ*=0.92, p<0.0001, Figure 5B).

**Fig. 5.**
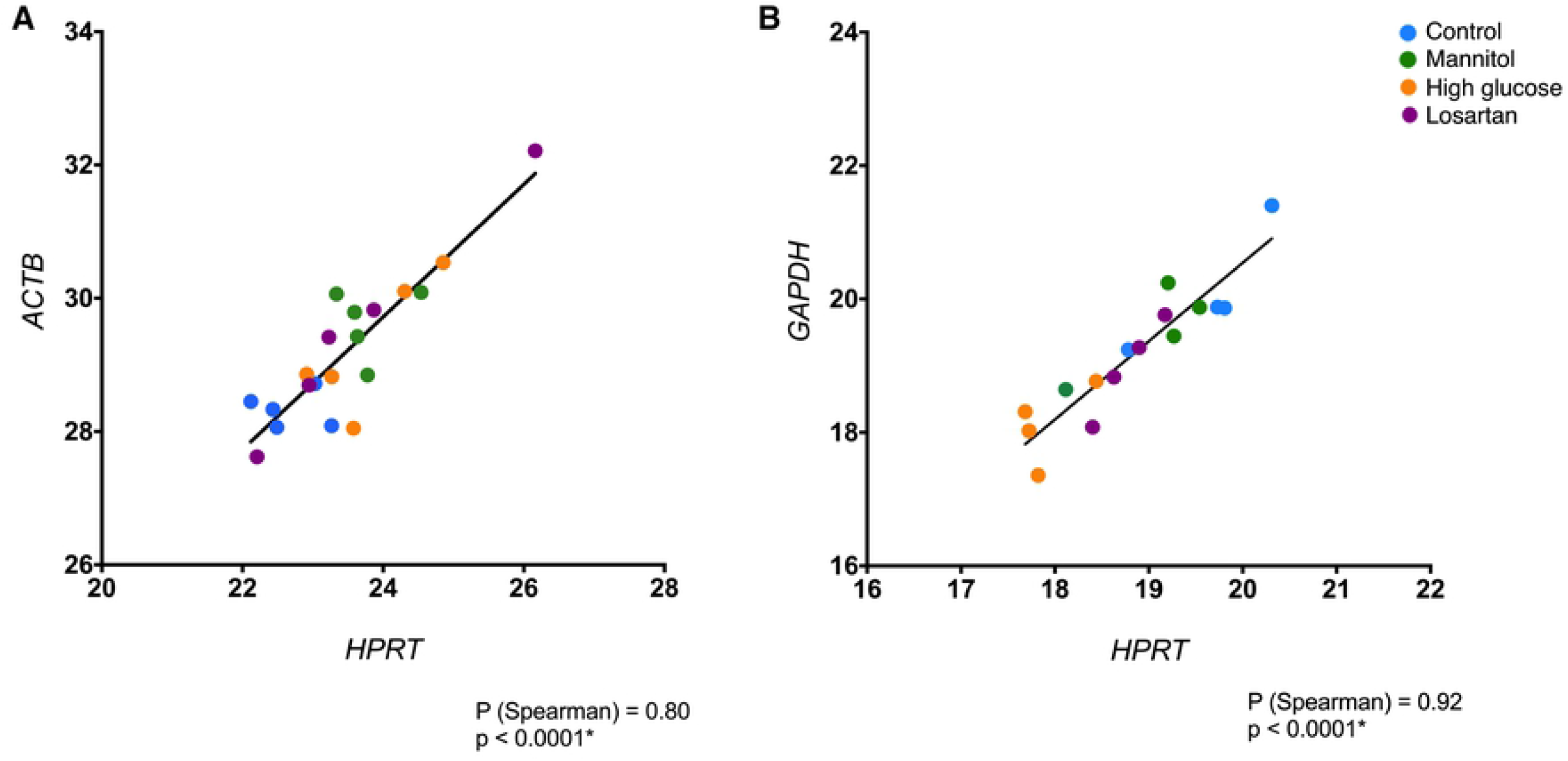
Correlation between top genes for both cell lines. A) Correlation between *ACTB* and *HPRT* expression profile in mesangial cells. B) Correlation between *GAPDH* and *HPRT* expression profile in podocytes. *ρ*: Spearman’s rank correlation coefficient. *p<0.0001.

### Validation of the best reference genes

As results showed that *HPRT* and *ACTB* were the best genes for normalization of mesangial cells qPCR data, we confirmed statistically that there was no difference among the four studied groups regarding the expression of these genes (p>0.05 by Kruskal-Wallis test) (Figure 3A). We also confirmed that there was no difference among the groups of podocytes regarding *HPRT* and *GAPDH* expression, the best genes for this cell type (Figure 3B). In fact, there was no difference between the groups for all candidate genes, in both cell lines.

## DISCUSSION

The pipeline used in this work has been extensively used throughout many laboratories and is accepted by literature as a reliable approach to determine the best reference gene to be used, specifically for qPCR in a predetermined biological sample and condition (18,20,21). Here, we aimed to provide data to determine the most suitable reference gene to be used for mesangial cells and podocytes exposed to a high glucose environment and treated with Losartan, a very known *in vitro* model for diabetic kidney disease (7,22–25).

Many research groups have clearly shown a need for studies that approach reference genes for their specific study sample (26,27). The importance of using the best-known reference gene and to pragmatically look at this question relies on the frequent inappropriate use of least feasible reference genes resulting in an inaccurate analysis of qPCR results, and therefore in loss of reagents, time, and samples. Sometimes the most known genes, such as *ACTB* and *GAPDH*, are used for samples and conditions that do not support their use. Even in the most recent years, researchers still normalize their qPCR data of *in vitro* studies based on the most frequently used genes, as *ACTB* for podocytes and *GAPDH* for mesangial cells (opposite to the finding we had in our analysis), without literature support for this choice (28–33). Unfortunately, some studies do not clearly provide which reference gene was used to normalize the data, or even if there was a data normalization. This shows the need of systematic analysis to identify the best gene or genes to be used as references.

In fact, literature frequently stands against the use of many popular reference genes. A systematic review performed on vertebrate studies found that 72% of the included studies used *GAPDH, ACTB* or *18S* as normalizing genes. The same group shows that as the number of screened reference genes for a specific study design increases, the chance of one of these three genes being the most stable decreases (34).

In nephrology, few studies have addressed reference genes for qPCR normalization (13), exposing a lack of information regarding which gene must be used for gene expression studies for kidney samples and cell lines. Kidney itself is an organ specifically characterized by numerous cell types, justifying the need for reference genes regarding each different cell line (35,36).

The genes selected as best ones for the studied samples - *HPRT* and *ACTB* for mesangial cells and *HPRT* along with *B2M* for podocytes - are extensively described in literature. Hypoxanthine phosphoribosyltransferase *(HPRT)* is mainly known by its role in the metabolism of purines, although impaired expression of this gene is also responsible for causing cell cycle dysregulation and multi-system regulatory dysfunction (37,38). Actin beta (*ACTB*) is involved in cell structure, motility, and integrity, and, as it is essential to multiple cell functions, the gene is highly abundant in many cell lines (39). Glyceraldehyde 3-phosphate dehydrogenase (*GAPDH*), although it is reported to be involved in cellular survival, apoptosis and DNA repair, is mainly known to express a cellular energy enzyme determinant to the glycolytic process, functioning as a catalyzer of triose phosphate oxidation and, for this reason, ubiquitously distributed in all cell types (40,41).

As long-established cell lines in literature, mesangial cells and podocytes are important biological samples to determine the best reference gene - many researchers in the field are focused on these structures (42–47) and the data provided by our work could potentially influence many studies, providing support to avoid wrong interpretation of results and its influence in downstream analysis and further conclusions.

## CONCLUSION

We analyzed six different genes using five software applications and the ΔCt method to determine that the best genes to be used for mesangial cell studies with high glucose and angiotensin receptor II blocker are *HPRT* and *ACTB,* while in the same conditions, the best combination of genes for podocyte gene expression normalization is *HPRT* together with *GAPDH*. We believe our work may provide support to many research laboratories engaged in mesangial cell and podocytes cell culture studies, allowing them to improve the quality of gene expression studies via qPCR and, consequently, the overall quality of nephrology research.

## Author contributions

NDH, ACA, and MAB designed the study. NDH was responsible for cell manipulation. NDH performed the experiments. NDH, ACA, and MAB analyzed the data. NDH, ACA, and MAB wrote the manuscript.

## Acknowledgements

We would like to thank Antonio S. Novaes for cell culture and qPCR technique training.

